# The pattern of mammal change in Garig Gunak Barlu (Cobourg) National Park suggests a fire-mediated pathway to decline and one for recovery

**DOI:** 10.64898/2026.05.21.726985

**Authors:** A.S. Kutt, A. Edwards, H.S. Fraser

**Author notes:** **Target:** *Wildlife* Research. **Funding:** This research did not receive any specific funding. **Data availability statement:** Data is available on request.

## Abstract

**Context:** Fire is a key driver of wildlife pattern in northern Australia, and in recent decades there has been concerted efforts to return burning practices from unmanaged wildfire to approaches more in tune with traditional approaches to fire management undertaken by First Nations people.

**Aims:** We examined fire pattern (early and late dry season) and distance from coast in relation to mammal pattern on a Cobourg peninsula, as there was some evidence in variable patterns in fire history and mammal persistence in this landscape.

**Methods:** In this study we examined data collected from Garig Gunak Barlu (Cobourg) National Park, Northern Territory, between 2004 and 2017. We investigated the relationship between all native mammal abundance and species richness, and abundance for eight species, with the three environmental variables (distance from coast, preceding early dry and late dry season burning frequency), using an information theoretic model selection and averaging framework.

**Key results:** There is a distinctive change in the fire and mammal pattern with respect to distance from the coast; that is, early and late dry season burning increasing moving inland from the coast, and a concomitant decrease or persistence in some key species on this fire gradient.

**Conclusions:** Our results suggest that fire management in northern Australia still needs targeted research linked to active management to more effectively prevent declines of mammal populations.

**Implications:** The location and shape of Garig Gunak Barlu (Cobourg) National Park provide an excellent opportunity to undertake some natural experiments in strategic burning to investigate improved approaches to conservation in fire-prone landscapes.

## Introduction

Fire is a dominant ecological pressure in tropical savannas, Australia and globally (Yates *et al*. 2008; Alvarado *et al*. 2020). In northern Australia, the shift from Aboriginal to European land management profoundly altered fire regimes. From the late 20^th^ century onwards, vast areas became dominated by uncontrolled, late dry season wildfires (Russell-Smith *et al*. 2003). In response, land managers and researchers developed tools and programs to reinstate traditional regimes based on early dry season prescribed burning (Price *et al*. 2012). This culminated in the establishment of a nationally accredited savanna burning emissions abatement methodology (Whitehead *et al*. 2014), enabling First Nations rangers and other groups to conduct strategic fire management while generating carbon credits (Russell-Smith *et al*. 2015). Long term programs have demonstrated that fire management can substantially reduce both the total area burnt and the extent of destructive late dry season wildfires (Ezzy 2022).

Beyond carbon accounting benefits, savanna burning programs are widely assumed to deliver biodiversity co-benefits, by reducing fire frequency, severity and extent. However, empirical evidence for these biodiversity gains remains patchy (Corey *et al*. 2020) with only some documented examples, such as in north-western Australia (Radford *et al*. 2020). Long term ecological monitoring at Kakadu National Park identified calamitous declines in small mammal populations from about 1996-2009 (Woinarski *et al*. 2010). Ongoing monitoring, and re-examination of the data from these sites, has concluded that this change in mammal abundance has not abated, and is correlated with fire size, fire frequency (Lawes *et al*. 2015) and distance from long-unburnt rugged and mesic vegetation (Einoder *et al*. 2023). Together, these findings highlight a pressing need to test whether savanna burning and carbon abatement programs can indeed deliver biodiversity benefits (Edwards *et al*. 2021).

In this study we examined data from Garig Gunak Barlu (Cobourg) National Park (GGB), Northern Territory, collected systematically between 2004 and 2017. The park is renowned for containing remnant populations of the Brush-tailed Rabbit Rat (*Conilurus penicillatus)*, though studies that modelled population persistence suggest ongoing declines in numbers regardless of fire regime and habitat type, and possible extirpation of the population in ten years (i.e., by 2020, Firth *et al*. 2010). We investigated mammal fauna trends in relation to fire history and location, particularly given GGB is a peninsula, and the pattern of change in mammals over time. We assessed change through correlation to fire history, (Edwards *et al*. 2003) distance from the coast and road-based ignitions. By exploring the spatial and temporal patterns of mammal decline with respect to fire, we aim to clarify whether fire management can support pathways to persistence and recovery.

## Methods

### Study Site

Data collection occurred in Garig Gunak Barlu National Park (formerly Gurig National Park and Cobourg Marine Park, and hereafter GGB) located 560 km northeast (-11° 19’ S, 132° 10’ E) of Darwin in the Northern Territory (NT) (Fig. 1). GGB occurs on a peninsula and is the northernmost site on mainland NT, with single road access only via Arnhem Land. GBB covers about 226,000 ha, the vegetation is predominantly Eucalypt dominated open forests and woodlands (tropical savanna), with patches of monsoon vine-thicket and closed forest, grasslands, wetlands and mangroves. The region is affected by the monsoon with distinct wet (Nov. to May) and dry (June to Oct.) seasons. Mean seasonal rainfall is 1,261 mm in the wet and 50 mm in the dry season, with a mean annual temperature of 31.5◦C (search Black Point NT, BoM). Whereby, annual fine fuel accumulation is abundant in the hot and humid wet season and subsequent high fire weather conditions lead to frequent and extensive area burnt in the dry season.

**Figure 1.**
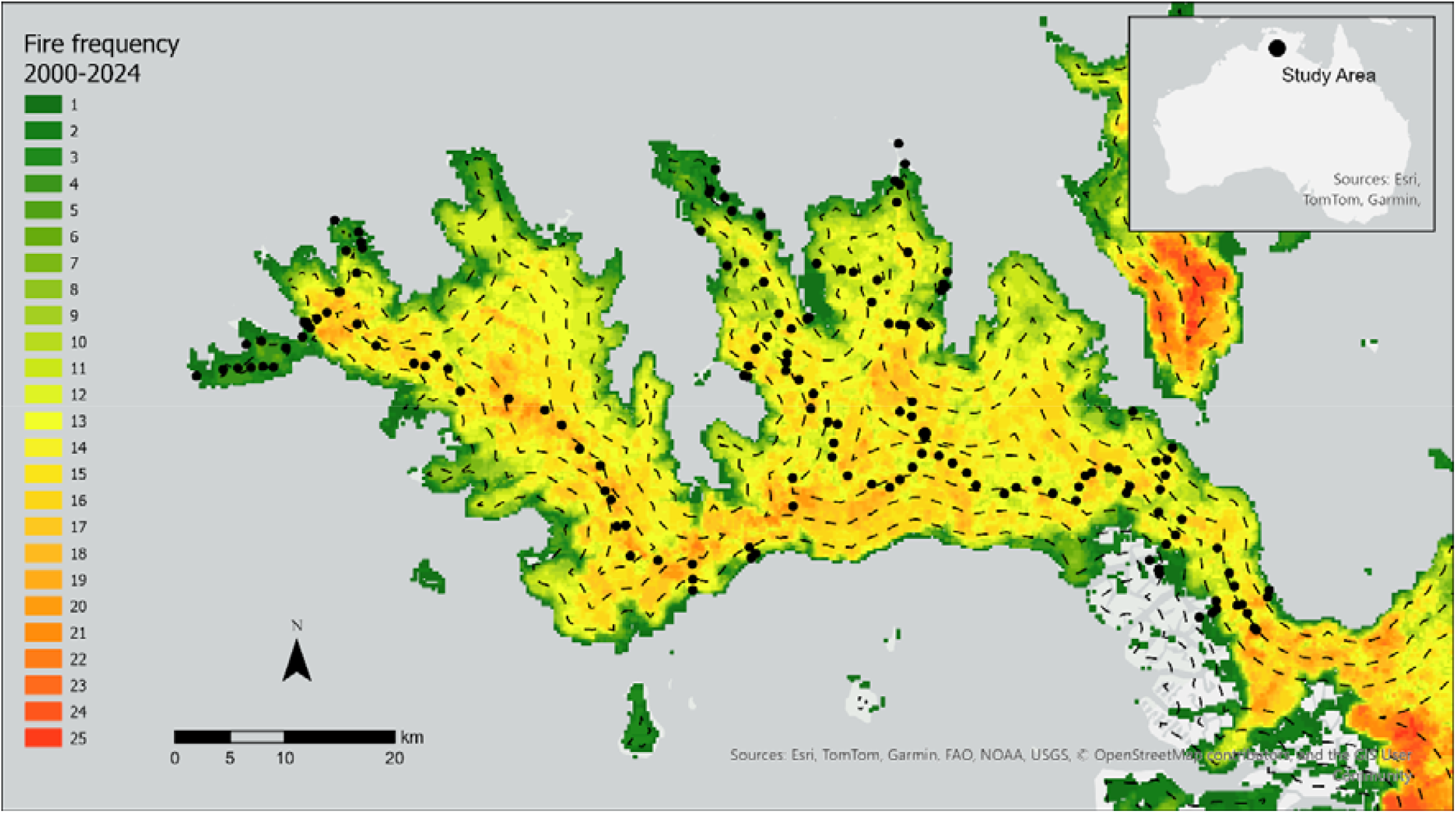
Garig Gunak Barlu (Cobourg) National Park, location and fire frequency mapping (2000-2024), with the distance from coast contours (dashed line) and survey sites (black circles) marked.

### Sampling

We examined data from seven surveys conducted at GGB between 2004 and 2017. Surveys were conducted largely to monitor changes in the mammal, bird and reptile fauna, though largely without any specific management question (i.e., the sites were unstratified with respect threats and management). The methods used were the systematic quadrat approaches undertaken throughout the Northern Territory (NTEPA 2013; Einoder *et al*. 2020). In this study we only considered the mammal data.

The standard methods (i.e., mammal records) and effort varied slightly across the sites and years. The number of trap nights ranged from two to four, the number of Elliott traps ranged from 16-30, the number of cages set ranged from 4-8, and the number of pitfall traps ranged from 0-4. Nocturnal active searching was consistent across all surveys (two 20-minute searches), though in the 2017 and 2018 surveys these were four 10-minute searches.

To address these variations, we standardised the data to represent species richness or abundance per 100 trap nights. The total trap effort was the sum of the Elliott, cage and pitfall traps set at each site multiplied by the number of nights of survey. Trap nights ranged from 72-108. The site species richness or abundance was divided by this figure and multiplied by 100. For example, a site sampled over 2 nights by 20 Elliotts, 4 cages and 4 pitfall traps, represents 56 trap nights. If a species scored an abundance of 1 for that site, then the relative abundance for 100 trap nights, is 1/(2*28) *100 or 1.78.

In 2011 the surveys only used Elliott and Cage traps, and active searching was not conducted in 2014. We address any potential limitations caused by this imbalance in the discussion.

### Environmental variables

The fire data were produced by North Australia and Rangelands Fire Information (NAFI) at Charles Darwin University, derived from satellite imagery sourced from the Moderate Resolution Imaging Spectroradiometer (MODIS) using the Red and Near Infrared bands with a 250 × 250 m pixel resolution. The fire metrics, frequency of early season burns (FRQE) and the frequency of late season burns (FRQL) were calculated using standard GIS techniques covering the 10 years prior to each of the survey years. Early dry season commences usually in May, at the end of the annual wet season (November to April), through to July, and the late dry season occurs from August to October, or later, depending on the commencement of the following wet season. The fire metrics were attributed to each site using the mean value of a 3 × 3-pixel window centred over the survey location. Distances from the coast were derived via standard GIS techniques, using ArcGIS Pro version 3.3.1 (ESRI Inc. 2024).

### Analysis

We only considered a subset of the mammal data recorded from the GGB surveys, due to the lack of records for many species. The subset included, all native mammals < 5.5 kg (nominal critical weight range), and recorded at > 10 sites from all surveys combined. The Agile Wallaby *Macropus agilis* was also included. Furthermore, we did not consider Northern Quoll *Dasyurus hallucatus*, as its decline is linked to invasion and subsequent poisoning by Cane Toads *Rhinella marinus*, and our study is focussed on fire responses. Lastly, we did not consider records of the Sugar Glider *Petaurus aeriel* as this species is largely recorded via spotlighting, though recognising species such as *Conilurus penicillatus* are also often recorded via spotlighting. We provide the mean abundance per site in each year, and for distance from coast classes for all these species in Supplementary Table 1 and 2.

As our contention in this study was that mammal species changed in abundance over time due to fire history, we initially undertook some simple exploratory analysis by plotting the mean (and standard error) of the mammal abundance and temporally appropriate fire metrics separated by distance to coast (i.e., < 1km, 1-3 km, 3-5 km, 5-8 km). We also plotted the proportion of late and early dry season fires per year from 2000-2024, which encompassed the survey years. We also undertook a univariate analysis of variance for native mammal abundance and species richness, to test year, distance classes and their interaction.

We investigated the relationship between all native mammal abundance and species richness, and abundance for eight species, with the three environmental variables, using an information–theoretic model selection and averaging framework. The environmental data were first standardised to a 0–1 range using a min–max transformation to allow direct comparison of the estimates. For each mammal variable, we fitted information-theoretical models with a negative binomial error distribution to account for overdispersion and the high frequency of zero counts. A global model including all three predictors was specified, and all possible subsets were generated using model dredging. Models were ranked according to the Akaike Information Criterion corrected for small sample size (AICc). Models with ΔAICc < 2 were retained as the best-supported candidate set, following the convention that such models are statistically indistinguishable in their level of empirical support (Burnham and Anderson 2002).

Where multiple candidate models were supported, we performed model averaging to estimate regression coefficients, unconditional standard errors, and 95% confidence intervals for each predictor. Akaike weights were used to calculate the relative support for each model, and the sum of weights across all models containing a predictor was taken as a measure of its relative importance. A high cumulative weight (e.g. ≥0.8) was interpreted as strong support that a predictor was influential in explaining variation in mammal abundance, whereas lower weights indicated weaker evidence for predictor effects. All analyses were conducted in R version 4.4.3 (R Core Team 2024). Negative binomial models were fitted using the *MASS* package (Venables and Ripley 2002) and model selection and averaging were carried out with the *MuMIn* package (Bartoń 2024). Other analysis were undertaking using Statistica (TIBCO Software Inc 2018).

## Results

A total of 15 native mammals were recorded from all surveys, included in the site species richness and abundance data. As indicated earlier, *Dasyurus hallucatus* and *Petaurus ariel* were not considered further in the analysis and also excluded were *Osphranter antilopinus* (only recorded at one site), *Sminthopsis nitela* (one site), *Trichosaurus vulpecula* (seven sites) *Phascogale tapoatafa* (one site) and *Pseudomys nanus* (eight sites) due to the low number of records.

The proportion of area unburnt, burnt in the early dry season and burnt in the late dry season, in each year, across the entire GGB National Park, altered from 2000 to 2024 (Fig. 2). From 2005, the total burnt area increased from 33% to 44%, the early dry season proportion burnt increased from 5% to 24% and late dry season proportion burnt decreased from 27% to 19% (Fig. 2). Though the early dry season area burnt increased, the total area burnt remained high. Late dry season burning declined in some years (2008-2014) but was also high in some years (2014-15, 2017-2019) (Fig. 2).

**Figure 2.**
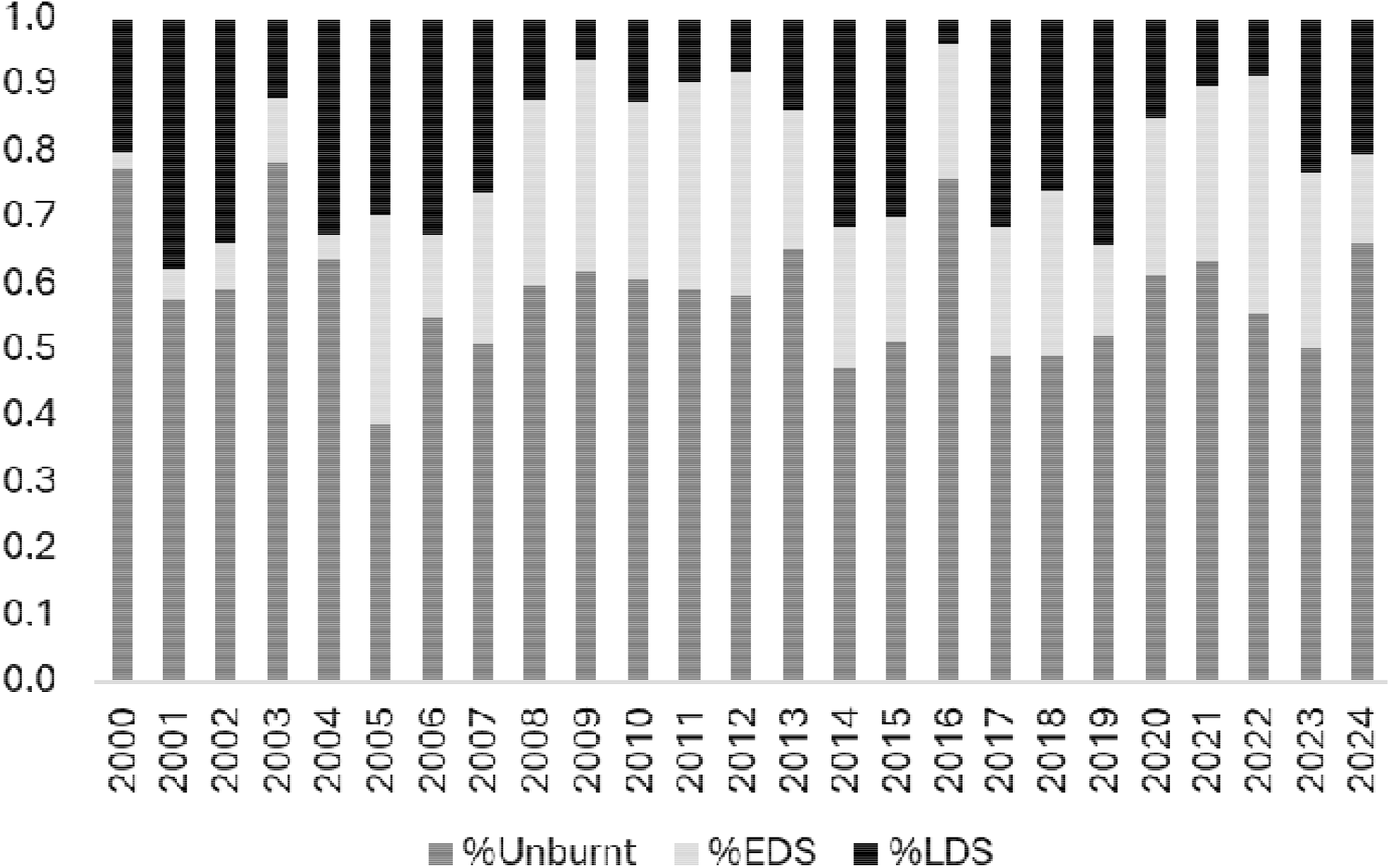
The proportion of unburnt, early dry season (%EDS) and late dry season (%LDS) burnt areas at Garig Gunak Barlu (Cobourg) National Park from 2000-2024.

Using fire data from the survey sites only, early and late dry season fire frequency changed significantly over the years of survey and was highest in 2014 and 2017 (Supplementary Table 1). Considering the four distances from coast categories, early and late dry season fire frequency was significantly higher in the 3-5 km and 5-8 km distance categories (Supplementary Table 2), which may correspond to the fire ignition locations. The pattern in the means for the two fire variables for each site, across the distance classes for each year, indicated a general pattern of increasing early and late fire frequency, the further the site was from the coast (Fig. 3).

**Figure 3.**
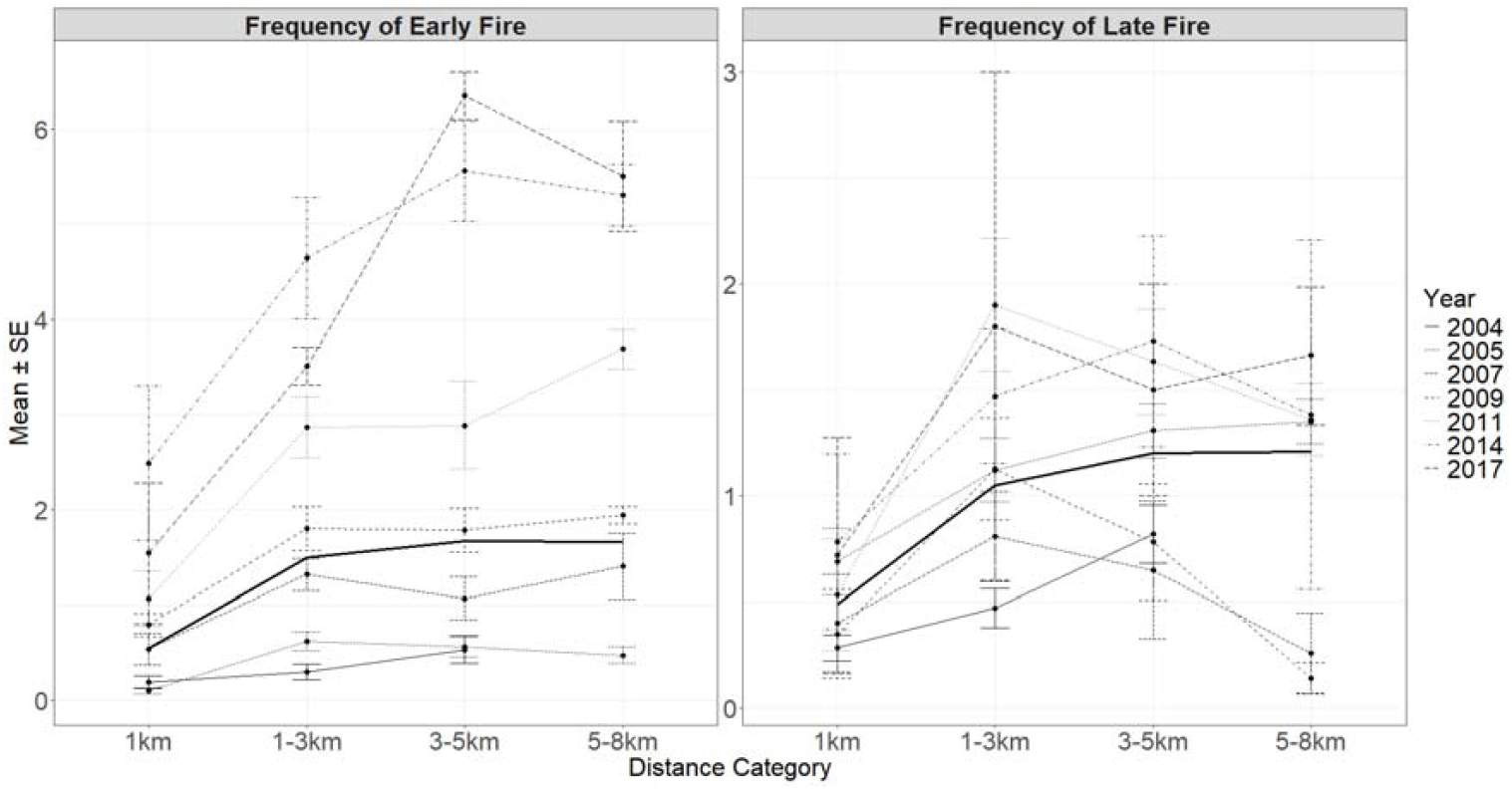
The mean (and standard error) of frequency of early and late dry season fires, separated by year of survey and distance from coast class. The black line is the mean for all survey years across the distance classes.

The univariate analysis of variance indicated that mammal abundance was significantly different in each year (df = 6, F=10.3, p <0.001), in each distance category (df = 3, F = 13.8, p = 0.024) and for the interaction or year and distance (df = 23, F = 1.7, p = 0.039). For mammal species richness, it was significantly different in each year (df = 6, F = 10.5, p <0.001), in each distance category (df = 3, F=9.1, p < 0.001) and not for the interaction (df = 23, F=1.3, p =0.231). The pattern in the means for the mammal groups and species for each site, across the distance classes for each year, indicated a decline the further the site was from the coast for *Antechinus bellus, Conilurus penicillatus, Macropus agilis, Melomys burtoni*, and *Pseudomys delicatulus* (Fig. 4). Conversely *Mesembriomys gouldii* and *Rattus tunneyi* were more abundant further from the coast. *Isoodon macrourus* demonstrated no correlation with distance to the coast (Fig. 4).

**Figure 4.**
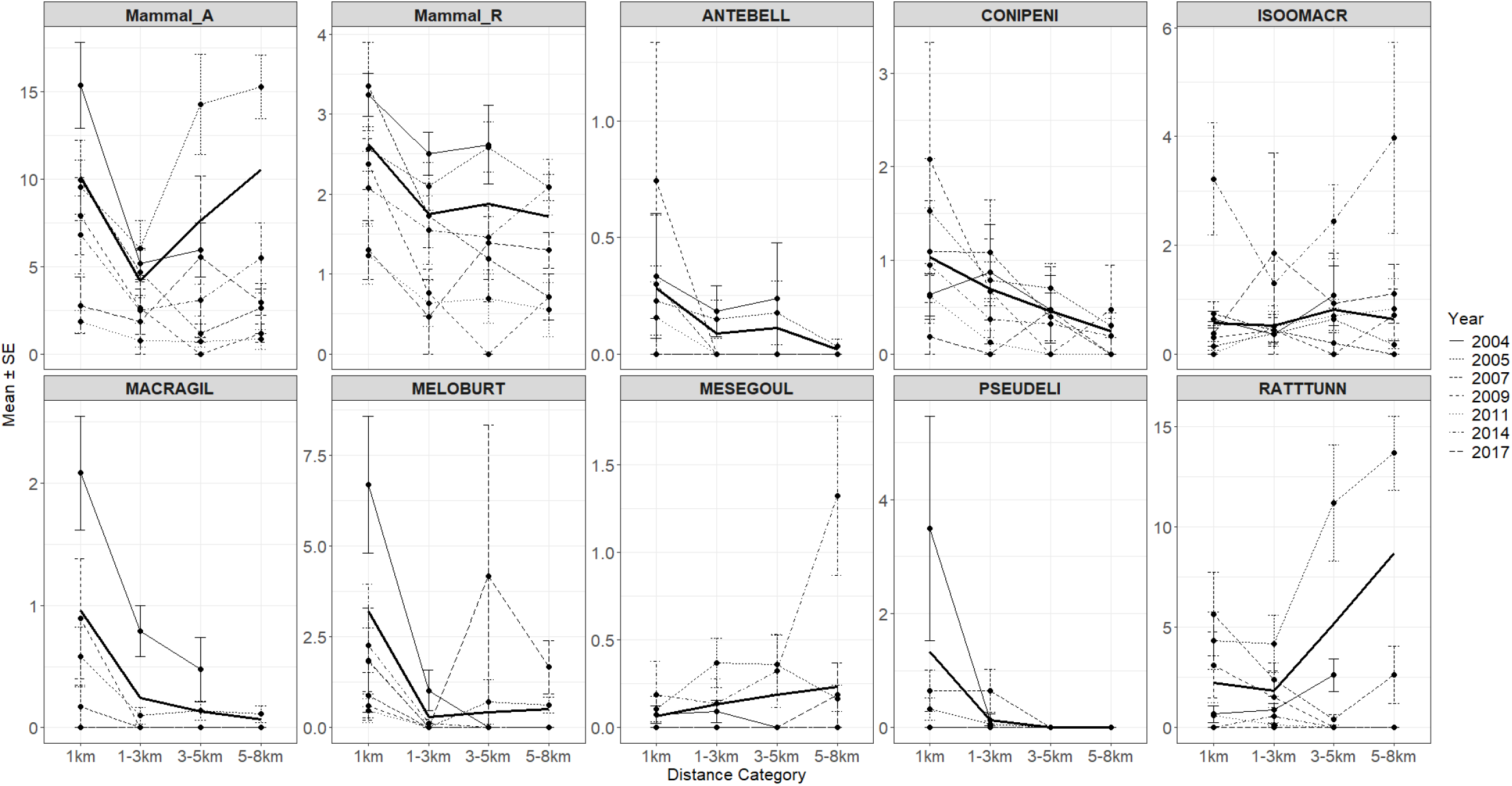
The variation in mammal abundance and mammal species richness (and standard error), and the abundance (and standard error) of eight mammal species separated by year of survey and distance from coast class. The black line is the mean for all survey years across the distance from coast classes.

Distance from coast was included in nine of the ten best supported AIC models, early fire frequency in seven of the best supported models and late fire frequency in two of the best supported models (Table 1). The model averaged estimates that were significant, indicated that there was a negative effect for distance (i.e., a decline the further the site was from the coast) for mammal species richness, *Macropus agilis, Conilurus penicillatus* and *Melomys burtoni* abundance, and a positive relationship for *Rattus tunneyi* and *Mesembriomys gouldii* abundance (Fig. 5). There was also a negative effect for frequency of early dry season fires (i.e., a decline with a higher frequency) for mammal abundance, mammal species richness, *Macropus agilis, Antechinus bellus, Conilurus penicillatus* and *Rattus tunneyi*, and a positive relationship (i.e., increase in abundance) for *Isoodon macroura* (Fig. 5). There was no significant effect identified by the model averaging for late season fire frequency.

**Table 1.** The two best supported AIC models for the mammal groups. K is the number of model terms (including the intercept and the negative-binomial dispersion parameter). LogLik is the log likelihood, AICc is the Akaike Information Criterion corrected for small sample size, and Delta AIC is the relative difference between the AIC of one model and the best (lowest) AIC in all the models tested.

| Group / Species | Model | K | LogLik | AICc | DeltaAIC |
| --- | --- | --- | --- | --- | --- |
| Mammal abundance | ~ Distance + FRQ_Early + 1 | 4 | -979.17 | 1966.46 | 0.00 |
| Mammal abundance | ~ Distance + FRQ_Early + FRQ_Late + 1 | 5 | -978.75 | 1967.68 | 1.22 |
| Mammal richness | ~ Distance + FRQ_Early + 1 | 4 | -552.33 | 1112.78 | 0.00 |
| Mammal richness | ~ Distance + FRQ_Early + FRQ_Late + 1 | 5 | -552.26 | 1114.70 | 1.92 |
| *Antechinus bellus* | ~ Distance + FRQ_Early + 1 | 4 | -91.71 | 191.54 | 0.00 |
| *Antechinus bellus* | ~ FRQ_Early + FRQ_Late + 1 | 4 | -92.29 | 192.72 | 1.17 |
| *Conilurus penicillatus* | ~ Distance + FRQ_Early + 1 | 4 | -300.41 | 608.94 | 0.00 |
| *Conilurus penicillatus* | ~ Distance + FRQ_Early + FRQ_Late + 1 | 5 | -299.46 | 609.11 | 0.17 |
| *Isoodon macroura* | ~ FRQ_Early + 1 | 3 | -307.98 | 622.03 | 0.00 |
| *Isoodon macroura* | ~ FRQ_Early + FRQ_Late + 1 | 4 | -307.59 | 623.31 | 1.28 |
| *Macropus agilis* | ~ Distance + FRQ_Early + 1 | 4 | -190.95 | 390.03 | 0.00 |
| *Macropus agilis* | ~ Distance + FRQ_Early + FRQ_Late + 1 | 5 | -190.90 | 392.00 | 1.96 |
| *Melomys burtoni* | ~ Distance + 1 | 3 | -308.49 | 623.05 | 0.00 |
| *Melomys burtoni* | ~ Distance + FRQ_Early + 1 | 4 | -308.18 | 624.48 | 1.43 |
| *Mesembriomys gouldii* | ~ Distance + 1 | 3 | -119.77 | 245.62 | 0.00 |
| *Mesembriomys gouldii* | ~ Distance + FRQ_Late + 1 | 4 | -119.47 | 247.06 | 1.44 |
| *Pseudomys delicatulus* | ~ Distance + FRQ_Late + 1 | 4 | -96.41 | 200.94 | 0.00 |
| *Pseudomys delicatulus* | ~ Distance + FRQ_Early + FRQ_Late + 1 | 5 | -95.61 | 201.41 | 0.47 |
| *Rattus tunneyi* | ~ Distance + FRQ_Early + FRQ_Late + 1 | 5 | -611.89 | 1233.97 | 0.00 |
| *Rattus tunneyi* | ~ Distance + FRQ_Early + 1 | 4 | -613.22 | 1234.56 | 0.59 |

**Figure 5.**
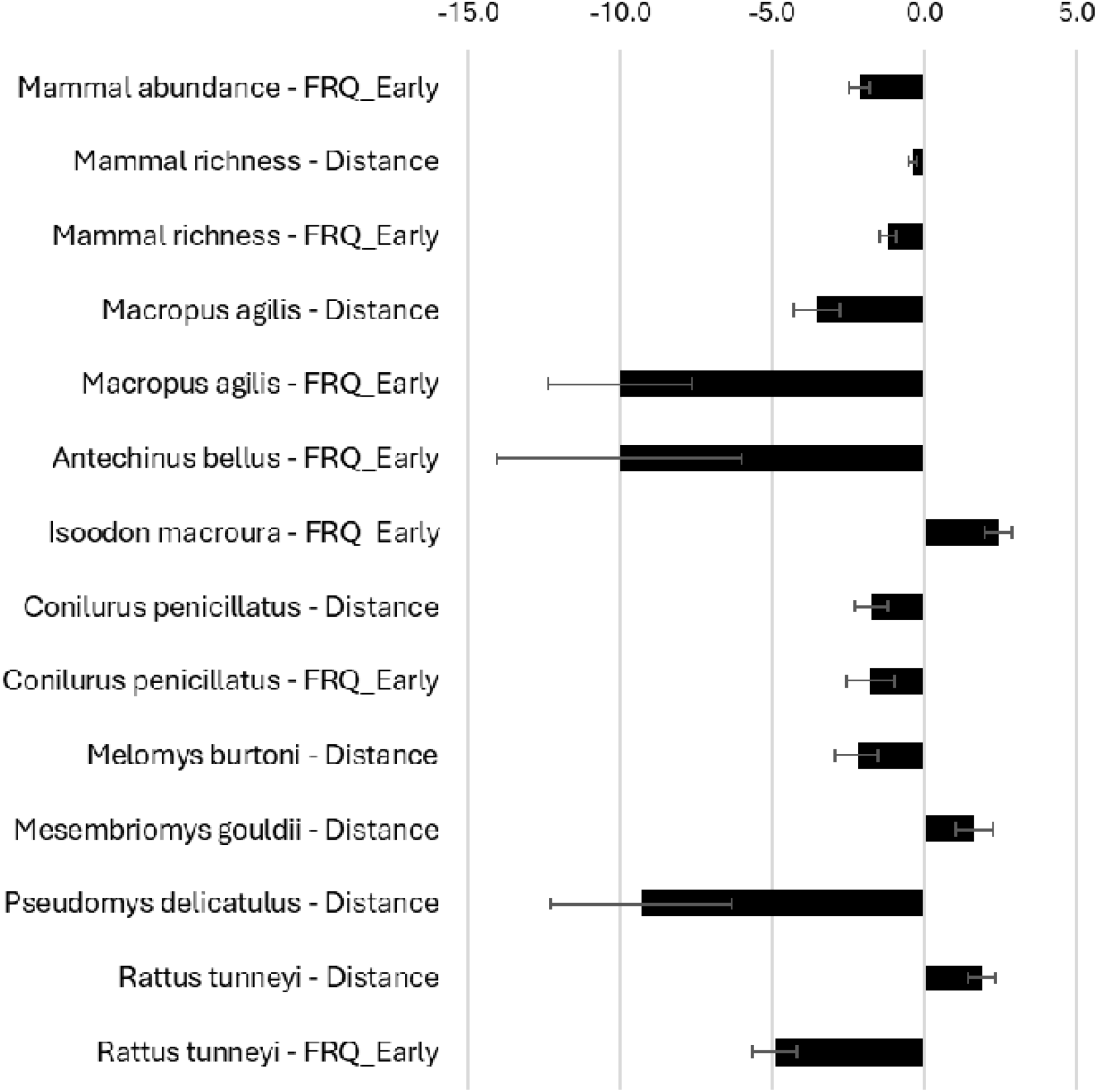
The significant parameter estimates and standard error for the mammal groups identified via model averaging. FRQ_Early is frequency of early dry season fires. Distance is the distance <u>from</u> the coast.

## Discussion

In this study we found that there was a change in the mammal fauna from the initial sampling in 2005 to the most recent sampling in 2018. This change was largely a decline for most species, and this is in keeping with other analysis of mammal fauna in the Top End of the Northern Territory, both in the past (Woinarski *et al*. 2010) and more recently (Neave *et al*. 2024). There has been a consistent link between changes in mammal population and fire (Lawes *et al*. 2015) and the reduction of long unburnt vegetation (Einoder *et al*. 2023), though some species are less affected by fire or even increase as the frequency of fire increases (Davies *et al*. 2022). This is often due to the presence or interaction with other threats such as feral cats *Felis catus* (Neave *et al*. 2024). Our data adds to the discourse, but present some important differences, notably, that there is a landscape pattern in the changes to mammals and fire, which point to both a cause and a solution.

We identify three key limitations in the interpretation of our data. Firstly, the association between the changes in mammal abundance and fire, is correlative. The initial and subsequent surveys were designed to provide surveillance data of the fauna in GGB and simply examine changes over time, without a clear management question. In addition there was no consistent site based habitat data collected in each year of survey that could be linked to biology and ecology of species, though detailed work was conducted on *Conilurus penicillatus* (Firth *et al*. 2010). There is also no consistent data regarding other threats that could affect the mammal populations, such as feral predators and herbivores which interact with fire to cumulatively impact on mammals. In addition, the site selection, though attempting to cover the range of habitats across the peninsula, largely followed the roads. The second limitation is the change in the number of sites sampled over time. Though the initial surveys in 2005 were focussed on discovery, the effort (i.e., number of sites) declined dramatically, which reduced the environmental variability of sites sampled. Lastly, there were subtle changes in the methods over time, and though we attempted to standardise our data, this might have had some impact on the relative abundance data and modelling, though we did try to discount species that might have been most affected by this (i.e., arboreal mammals).

Notwithstanding these limitations, the data indicated a significant change in mammal species richness and abundance at each site over the years of survey. Some changes, such as the loss of *Dasyurus hallucatus*, are due to the national-scale waves of quoll decline due to the east-west invasion of Cane Toad *Rhinella marinus* (Moore *et al*. 2022). Similarly, the Pale Field Rat *Rattus tunneyi* seems to have disappeared across most of its range, without any confident cause (Braithwaite and Griffiths 1996; Tuft *et al*. 2021), though the more disturbance tolerant Delicate Mouse *Pseudomys delicatulus* (Kutt and Woinarski 2007) remained unrecorded after 2007. Some species were uncommon and idiosyncratic in pattern (e.g., Black-footed Tree-rat *Mesembriomys gouldii*), however there is an overall decline across the years of survey, as well as a concurrent increase in late and early dry season fires.

There is a distinctive change in the fire and mammal pattern with respect to distance from the coast. At a granular scale, the pattern of fire frequency for the entire peninsula indicates that there is a noticeable gradient with increasing distance from the coast (Fig. 1); the site data also indicated an increasing mean early and late dry fire frequency the further a site was from the coast, a pattern that consolidated over the subsequent years of survey. Though there is a strong imperative to shift annual burning to reduce late dry season fires (Ezzy 2022), our data indicated for the GGB survey sites, both early and late dry season burning increased, and did so more in the central portions of the national park, possibly associated with fires being ignited from the roads. At the park scale, the area of unburnt vegetation decreased overall. It is apparent that the goal of reducing fire frequency overall has not been achieved, whether due to management choices or climate factors. In high rainfall regions such as GGB, the suitable conditions for burning occurs later in the year (i.e., July to October, Perry *et al*. 2020) suggesting early dry season fires (i.e., before July) may be more difficult to achieve, if curing occurs later and the ability to conduct burning is reduced (i.e., fires will occur more readily in the late dry season). The fire data suggests that early dry season fires do not commence until July (Andrew Edwards, pers. obs.) and this could be due to access constraints after the wet season.

Our modelling indicated that mammals were on the whole declining each year and with increasing distance from the coast, apart from *Isoodon macroura* (consistent numbers over years and distance) and *Rattus tunneyi* (more abundant away from the coast, but declining year on year), a pattern for this rodent apparent for many decades elsewhere (Braithwaite and Griffiths 1996). Though vulnerable to frequent fires, which affect its survival (Pardon *et al*. 2003), *I. macroura* is also recorded as disturbance tolerant, though this may be a function of the presence of other threats (Davies *et al*. 2022). What was notable was the number of species that significantly declined in abundance both with increasing early dry season fire and/or distance from coast (which could be a function of both the habitat change, and fire frequency). This is consistent with the literature which shows that in Kakadu, which is adjacent to GBB, *Melomys burtoni* populations were projected to decline with increasing fire frequency (Griffiths *et al*. 2015) and *Antechinus bellus* is most abundant in unburnt sub-catchments (Andersen 2021). Similarly, total abundance and species richness of small mammals, including a majority of the common small mammals, have all been found to be affected by fire treatments (Corbett *et al*. 2003). Our data suggests that fire overall (and threats that interact with fire) and the location in the landscape, is having an ongoing negative effect on the mammal fauna.

In conclusion, our study indicated that there seems to be an approach to intentional or unintentional burning at GGB that is resulting in ongoing negative change to the mammals. Part of this may be due to the park’s location in the wetter northern tip of the Northern Territory (and therefore a naturally later fire season), the shape of the park as a peninsula (and the associated high extent of coastal vegetation) and the limited central road infrastructure. We reiterate that there are some limitations to our data (i.e., reduction in sites surveyed over time, and associated reduction in correlative inference) that may influence our results; however some simple patterns are apparent, that have potentially important and positive implications for fire management and mammal conservation into the future. Firstly, even though many of the late season fires that are occurring at GBB may be due to unintended fires caused by visiting tourists, the geographical location, and natural tendency for later fires to occur due to regional climate patterns (Perry *et al*. 2020), suggests the emphasis on undertaking early dry season fires should be increased, even if conditions are still wet. Small patchier strategic burns conducted earlier and aerially, could be one solution. Secondly, because of the shape of GBB with multiple fingers and smaller peninsulas (Fig. 1) reducing or eliminating annual fire in key zones (i.e., via strategic burning, fire breaks, or limiting access) could improve mammal persistence or numbers. Overall, the shape of GGB provides an excellent opportunity to undertake strategic fire planning, and natural experiments, similar to that conducted at Kapalga in Kakadu National Park (Andersen *et al*. 2003), to help inform fire management. Progress on small mammal recovery in northern Australia, cannot occur without careful consideration of the landscapes, climate and the opportunity to adapt management to local circumstances and data, rather than follow formulas that may not be always relevant (Russell-Smith *et al*. 2015).

## Acknowledgements

We acknowledge and pay our respects to the Arrarrkbi people who jointly manage Garig Gunak Barlu National Park. The surveys were conducted by many staff and volunteers from the Flora and Fauna Division, Department of Lands, Planning and Environment. All surveys were undertaken under scientific research permits, and animal ethics approvals, as prescribed by the Territory Parks and Wildlife Conservation Act. Fire information was created and generously supplied by the North Australia and Rangelands Fire Information service at Charles Darwin University.

