## Supplemental Tables for "The pattern of mammal change in Garig Gunak Barlu (Cobourg) National Park suggests a fire-mediated pathway to decline and one for recovery"

**Supplementary Table 1.** The mean (and standard error) for the environmental variables and the mammal groups and species. H is the non-parametric analysis of variance (Kruskal-Wallis) statistic, and p is the significance level. ns is not significant. Bold data is the highest mean, irrespective of the significance level.

| **Variable/ group / species** | **2004** | **2005** | **2007** | **2009** | **2011** | **2014** | **2017** | **H** | **p** |
| --- | --- | --- | --- | --- | --- | --- | --- | --- | --- |
| Number of sites | 68 | 120 | 30 | 30 | 31 | 29 | 14 |  |  |
| Fire frequency - Early | 0.28 (0.05) | 0.44 (0.05) | 1.07 (0.12) | 1.55 (0.13) | 2.48 (0.24) | **4.55 (0.36)** | 3.92 (0.62) | 181.6 | <0.001 |
| Fire frequency - Late | 0.43 (0.05) | 1.15 (0.07) | 0.58 (0.12) | 0.69 (0.13) | 1.36 (0.17) | **1.37 (0.24)** | 1.32 (0.28) | 54.6 | <0.001 |
| Abundance | 10.09 (1.36) | **11.96 (1.07)** | 5.03 (0.99) | 3.26 (0.95) | 1.08 (0.26) | 4.15 (0.73) | 3.11 (0.84) | 56.5 | <0.001 |
| Species richness | **2.87 (0.18)** | 2.30 (0.13) | 1.88 (0.29) | 1.03 (0.21) | 0.81 (0.17) | 1.75 (0.21) | 1.19 (0.18) | 77.9 | <0.001 |
| *Macropus agilis* | **1.35 (0.25)** | 0.22 (0.06) | 0.05 (0.05) | 0.24 (0.14) |  |  |  | 83.8 | <0.001 |
| *Antechinus bellus* | **0.26 (0.14)** | 0.13 (0.05) | 0.08 (0.08) | 0.20 (0.16) | 0.04 (0.04) |  |  | 5.1 | ns |
| *Isoodon macroura* | 0.60 (0.13) | 0.31 (0.07) | 0.40 (0.13) | 0.36 (0.14) | 0.45 (0.15) | **2.53 (0.48)** | 0.93 (0.32) | 51.3 | <0.001 |
| *Dasyurus hallucatus* | **0.19 (0.13)** | 0.09 (0.03) |  |  |  |  |  |  | ns |
| *Petaurus aerial* | **0.14 (0.06)** | 0.03 (0.02) | 0.04 (0.04) |  |  |  | 0.13 (0.09) |  | ns |
| *Conilurus penicillatus* | 0.70 (0.23) | 0.76 (0.15) | 0.77 (0.26) | **0.88 (0.41)** | 0.22 (0.09) | 0.44 (0.17) | 0.13 (0.09) | 5.2 | ns |
| *Melomys burtoni* | **3.54 (0.98)** | 0.78 (0.26) | 0.24 (0.17) | 0.16 (0.11) | 0.13 (0.10) | 0.51 (0.37) | 1.85 (0.76) | 35.7 | <0.001 |
| *Mesembriomys gouldii* | 0.07 (0.03) | 0.24 (0.06) |  |  |  | **0.44 (0.14)** | 0.07 (0.07) | 25.6 | <0.001 |
| *Pseudomys delicatulus* | **1.70 (0.94)** | 0.08 (0.04) | 0.41 (0.17) |  |  |  |  | 21.1 | <0.01 |
| *Rattus tunneyi* | 1.04 (0.27) | **9.19 (1.08)** | 2.90 (0.73) | 1.39 (0.66) | 0.22 (0.18) | 0.20 (0.16) |  | 101.5 | <0.001 |

**Supplementary Table 2.** The mean (and standard error) for the environmental variables and the mammal groups and species, for each distance class. H is the non-parametric analysis of variance (Kruskal-Wallis) statistic, and p is the significance level. ns is not significant. Bold data is the highest mean, irrespective of the significance level.

| **Variable/ group / species** | **0-1 km** | **1-3 km** | **3-5 km** | **5-8 km** | **H** | **p** |
| --- | --- | --- | --- | --- | --- | --- |
| Number of sites | 94 | 95 | 64 | 69 |  |  |
| Fire frequency - Early | 0.55 (0.1) | 1.49 (0.17) | **1.66 (0.24)** | **1.66 (0.24)** | 36.6 | <0.001 |
| Fire frequency - Late | 0.49 (0.07) | 1.05 (0.09) | 1.20 (0.10) | **1.21 (0.11)** | 50.2 | <0.001 |
| Abundance | 10.15 (1.08) | 4.17 (0.55) | 7.64 (1.44) | **10.55 (1.37)** | 31.1 | <0.001 |
| Species richness | **2.62 (0.15)** | 1.75 (0.15) | 1.88 (0.2) | 1.72 (0.13) | 22.0 | <0.001 |
| *Macropus agilis* | **0.96 (0.2)** | 0.24 (0.07) | 0.13 (0.05) | 0.07 (0.04) | 31.9 | <0.001 |
| *Antechinus bellus* | **0.28 (0.11)** | 0.09 (0.04) | 0.11 (0.07) | 0.02 (0.02) | 5.0 | ns |
| *Isoodon macroura* | 0.57 (0.12) | 0.53 (0.1) | **0.82 (0.16)** | 0.64 (0.21) | 2.7 | ns |
| *Dasyurus hallucatus* | **0.15 (0.09)** | 0.04 (0.03) | 0.10 (0.05) |  |  | ns |
| *Petaurus aerial* | 0.05 (0.03) | 0.04 (0.02) | **0.11 (0.05)** | 0.02 (0.02) |  | ns |
| *Conilurus penicillatus* | **1.04 (0.21)** | 0.69 (0.17) | 0.46 (0.13) | 0.24 (0.09) | 12.4 | <0.01 |
| *Melomys burtoni* | **3.20 (0.74)** | 0.29 (0.16) | 0.43 (0.29) | 0.51 (0.16) | 43.8 | <0.001 |
| *Mesembriomys gouldii* | 0.07 (0.03) | 0.13 (0.04) | 0.19 (0.08) | **0.23 (0.07)** | 4.0 | ns |
| *Pseudomys delicatulus* | **1.34 (0.69)** | 0.13 (0.06) |  |  | 17.1 | <0.001 |
| *Rattus tunneyi* | 2.23 (0.51) | 1.82 (0.43) | 5.17 (1.38) | **8.71 (1.39)** | 33.8 | <0.001 |

**Supplementary Table 3.** The results of the model averaging indicating the group, predictor, the estimate regression coefficients, unconditional standard errors, and 95% confidence intervals for each predictor. Akaike weights were used to calculate the relative support for each model, and the sum of weights across all models containing a predictor was taken as a measure of its relative importance. A high cumulative weight (e.g. ≥0.8) was interpreted as strong support that a predictor was influential in explaining variation in mammal abundance, whereas lower weights indicated weaker evidence for predictor effects.

| **Group/Species** | **Predictor** | **Estimate** | **StdError** | **LowerCI** | **UpperCI** | **ModelWeight** | **CumulativeWeight** |
| --- | --- | --- | --- | --- | --- | --- | --- |
| Abundance | Distance | 0.42 | 0.23 | -0.04 | 0.87 | 1.00 | 1.00 |
| Abundance | FRQ_Early | -2.13 | 0.34 | -2.80 | -1.46 | 1.00 | 2.00 |
| Abundance | FRQ_Late | 0.14 | 0.32 | -0.49 | 0.78 | 0.35 | 2.35 |
| Abundance | (Intercept) | 2.15 | 0.11 | 1.93 | 2.37 | 0.00 | 2.35 |
| Richness | Distance | -0.37 | 0.15 | -0.67 | -0.07 | 1.00 | 1.00 |
| Richness | FRQ_Early | -1.19 | 0.26 | -1.69 | -0.68 | 1.00 | 2.00 |
| Richness | FRQ_Late | -0.03 | 0.16 | -0.35 | 0.29 | 0.28 | 2.28 |
| Richness | (Intercept) | 0.96 | 0.07 | 0.83 | 1.10 | 0.00 | 2.28 |
| *Macropus agilis* | Distance | -3.52 | 0.74 | -4.98 | -2.07 | 1.00 | 1.00 |
| *Macropus agilis* | FRQ_Early | -10.01 | 2.36 | -14.66 | -5.37 | 1.00 | 2.00 |
| *Macropus agilis* | FRQ_Late | -0.12 | 0.69 | -1.47 | 1.23 | 0.27 | 2.27 |
| *Macropus agilis* | (Intercept) | 0.47 | 0.23 | 0.00 | 0.93 | 0.00 | 2.27 |
| *Antechinus bellus* | FRQ_Early | -10.03 | 4.02 | -17.94 | -2.12 | 1.00 | 1.00 |
| *Antechinus bellus* | Distance | -1.07 | 1.25 | -3.53 | 1.39 | 0.59 | 1.59 |
| *Antechinus bellus* | FRQ_Late | -1.11 | 2.01 | -5.06 | 2.84 | 0.41 | 2.00 |
| *Antechinus bellus* | (Intercept) | -0.92 | 0.51 | -1.91 | 0.08 | 0.00 | 2.00 |
| *Isoodon macroura* | FRQ_Early | 2.47 | 0.45 | 1.59 | 3.35 | 1.00 | 1.00 |
| *Isoodon macroura* | FRQ_Late | -0.14 | 0.42 | -0.97 | 0.68 | 0.26 | 1.26 |
| *Isoodon macroura* | Distance | -0.07 | 0.22 | -0.51 | 0.37 | 0.24 | 1.50 |
| *Isoodon macroura* | (Intercept) | -1.08 | 0.17 | -1.41 | -0.76 | 0.00 | 1.50 |
| *Conilurus penicillatus* | Distance | -1.74 | 0.54 | -2.80 | -0.68 | 1.00 | 1.00 |
| *Conilurus penicillatus* | FRQ_Early | -1.76 | 0.80 | -3.34 | -0.19 | 1.00 | 2.00 |
| *Conilurus penicillatus* | FRQ_Late | 0.60 | 0.90 | -1.17 | 2.37 | 0.48 | 2.48 |
| *Conilurus penicillatus* | (Intercept) | 0.15 | 0.24 | -0.33 | 0.62 | 0.00 | 2.48 |
| *Melomys burtoni* | Distance | -2.21 | 0.71 | -3.61 | -0.81 | 1.00 | 1.00 |
| *Melomys burtoni* | FRQ_Early | -0.17 | 0.58 | -1.31 | 0.97 | 0.26 | 1.26 |
| *Melomys burtoni* | FRQ_Late | -0.15 | 0.66 | -1.45 | 1.14 | 0.21 | 1.47 |
| *Melomys burtoni* | (Intercept) | 0.75 | 0.33 | 0.11 | 1.39 | 0.00 | 1.47 |
| *Mesembriomys gouldii* | Distance | 1.65 | 0.62 | 0.44 | 2.87 | 1.00 | 1.00 |
| *Mesembriomys gouldii* | FRQ_Late | -0.24 | 0.74 | -1.70 | 1.22 | 0.25 | 1.25 |
| *Mesembriomys gouldii* | FRQ_Early | 0.13 | 0.45 | -0.76 | 1.02 | 0.23 | 1.48 |
| *Mesembriomys gouldii* | (Intercept) | -2.80 | 0.37 | -3.52 | -2.08 | 0.00 | 1.48 |
| *Pseudomys delicatulus* | Distance | -9.31 | 2.97 | -15.16 | -3.46 | 1.00 | 1.00 |
| *Pseudomys delicatulus* | FRQ_Late | -7.90 | 5.09 | -17.89 | 2.10 | 0.83 | 1.83 |
| *Pseudomys delicatulus* | FRQ_Early | -3.56 | 4.63 | -12.65 | 5.52 | 0.54 | 2.36 |
| *Pseudomys delicatulus* | (Intercept) | 1.29 | 0.61 | 0.09 | 2.49 | 0.00 | 2.36 |
| *Rattus tunneyi* | Distance | 1.90 | 0.44 | 1.04 | 2.76 | 1.00 | 1.00 |
| *Rattus tunneyi* | FRQ_Early | -4.91 | 0.73 | -6.35 | -3.48 | 1.00 | 2.00 |
| *Rattus tunneyi* | FRQ_Late | 0.83 | 0.95 | -1.03 | 2.69 | 0.57 | 2.57 |
| *Rattus tunneyi* | (Intercept) | 0.87 | 0.25 | 0.39 | 1.36 | 0.00 | 2.57 |

**Supplementary Table 4.** The survey effort per year including the number of sites sampled.

| **Year** | **Sites** | **Nights** | **Elliotts** | **Cages** | **Pitfall** | **Active Search** |
| --- | --- | --- | --- | --- | --- | --- |
| 2004 | 68 | 3 | 20 | 4 | 4 | 5 x 20 min |
| 2005 | 134 | 3 | 20 | 4 | 4 | 5 x 20 min |
| 2007 | 30 | 3 | 20 | 4 | 4 | 5 x 20 min |
| 2009 | 30 | 3 | 20 | 4 | 4 | 5 x 20 min |
| 2011 | 31 | 3 | 20 | 4 | 0 | 0 |
| 2014 | 29 | 4 | 16 | 2 | 4 | 0 |
| 2017 | 14 | 4 | 16 | 8 | 3 | 10 x 10 min |
